# Perturbations in mitochondrial metabolism associated with defective cardiolipin biosynthesis: An *in-organello* real-time NMR study

**DOI:** 10.1101/2024.06.18.599628

**Authors:** Antonio J. Rua, Wayne Mitchell, Steven M. Claypool, Nathan N. Alder, Andrei T. Alexandrescu

**Affiliations:** Department of Molecular and Cellular Biology, University of Connecticut, Storrs, CT 06269, USA; Department of Physiology, Johns Hopkins University School of Medicine, Baltimore, MD 21205, USA; Mitochondrial Phospholipid Research Center, Johns Hopkins University School of Medicine, Baltimore, MD 21205, USA; Department of Genetic Medicine, Johns Hopkins University School of Medicine, Baltimore, MD 21205, USA

**Keywords:** metabolic disease, nuclear magnetic resonance (NMR), tricarboxylic acid (TCA) cycle, Krebs cycle, adenosine triphosphate (ATP), 3-methylglutaconic acid (3MGA), Barth syndrome (BTHS), mitochondrial respiration

## Abstract

Mitochondria are central to cellular metabolism; hence, their dysfunction contributes to a wide array of human diseases including cancer, cardiopathy, neurodegeneration, and heritable pathologies such as Barth syndrome. Cardiolipin, the signature phospholipid of the mitochondrion promotes proper cristae morphology, bioenergetic functions, and directly affects metabolic reactions carried out in mitochondrial membranes. To match tissue-specific metabolic demands, cardiolipin typically undergoes an acyl tail remodeling process with the final step carried out by the phospholipid-lysophospholipid transacylase tafazzin. Mutations in the *tafazzin* gene are the primary cause of Barth syndrome. Here, we investigated how defects in cardiolipin biosynthesis and remodeling impact metabolic flux through the tricarboxylic acid cycle and associated pathways in yeast. Nuclear magnetic resonance was used to monitor in real-time the metabolic fate of ^13^C_3_-pyruvate in isolated mitochondria from three isogenic yeast strains. We compared mitochondria from a wild-type strain to mitochondria from a Δ*taz1* strain that lacks tafazzin and contains lower amounts of unremodeled cardiolipin, and mitochondria from a Δ*crd1* strain that lacks cardiolipin synthase and cannot synthesize cardiolipin. We found that the ^13^C-label from the pyruvate substrate was distributed through about twelve metabolites. Several of the identified metabolites were specific to yeast pathways, including branched chain amino acids and fusel alcohol synthesis. Most metabolites showed similar kinetics amongst the different strains but mevalonate and α-ketoglutarate, as well as the NAD+/NADH couple measured in separate nuclear magnetic resonance experiments, showed pronounced differences. Taken together, the results show that cardiolipin remodeling influences pyruvate metabolism, tricarboxylic acid cycle flux, and the levels of mitochondrial nucleotides.

---

Mitochondria are the metabolic powerhouses of eukaryotic cells ^1-3^, typically generating >90% of the cell’s adenosine triphosphate (ATP) ^4^. Besides their roles in energy metabolism, mitochondria also coordinate lipid and nucleic acid biosynthesis, ion homeostasis, apoptosis, and inflammatory signaling ^3,5^. Because of their central role in metabolism, mitochondrial dysfunction is associated with a diverse array of human diseases that are caused by metabolic alterations ^6-8^. These include cancer, heart disease, neurodegenerative conditions, and heritable (primary) mitochondrial pathologies ^9,10^ such as Barth syndrome (BTHS) ^11,12^ and dilated cardiomyopathy with ataxia (DCMA) ^8^. Importantly, conditions such as BTHS and DCMA are directly associated with defects in the biosynthesis of cardiolipin (CL), the signature phospholipid of the mitochondrion^8,13^.

CL is a lipid with two phosphate headgroups and four acyl tails, predominantly located in the inner mitochondrial membrane (IMM) where it accounts for ∼10-20 mol% of membrane lipids ^14,15^. CL has an inverted conical molecular geometry that can promote bilayer curvature, which may be important for proper mitochondrial cristae formation and activity of membrane-associated mitochondrial enzymes ^16-18^. Following *de novo* synthesis in the IMM, precursor CL (pCL) undergoes de- and re-acylation steps termed ‘remodeling’ to yield mature CL (mCL) with a specific acyl composition (Fig. 1A). Disruptions in CL remodeling can impair mitochondrial metabolism, as several tissues synthesize CL with specific acyl chain profiles to meet their energetic requirements ^12,19^. Tafazzin (encoded by the *TAZ*/*TAZ1* genes in mammals and yeast, respectively) is an intermembrane space (IMS)-facing membrane-bound phospholipid transacylase that carries out the acyl transfer step in CL remodeling, transferring an acyl chain from a donor lipid to monolyso-cardiolipin (MLCL) to create the mature tetra-acyl CL ^14,20^. Mutations in the *TAZ* gene are the cause of the rare X-linked genetic disease Barth syndrome (BTHS) that affects about one in 400,000 males ^11,12,21,22^. BTHS is characterized by metabolic perturbations, diminished growth, fatigue, neutropenia, skeletal abnormalities, and cardiomyopathy, resulting in reduced quality of life and life expectancy ^22,23^. At the molecular level, BTHS patients show elevated MLCL:CL ratios, increased levels of the metabolite 3MGA (3-methylglutaconic acid) in plasma and urine, hypocholesterolemia, and mitochondrial dysfunction including altered morphology, increased reactive oxygen species (ROS) production, decreased ATP synthesis, and impaired oxidative phosphorylation ^22,23^.

**Figure 1:**
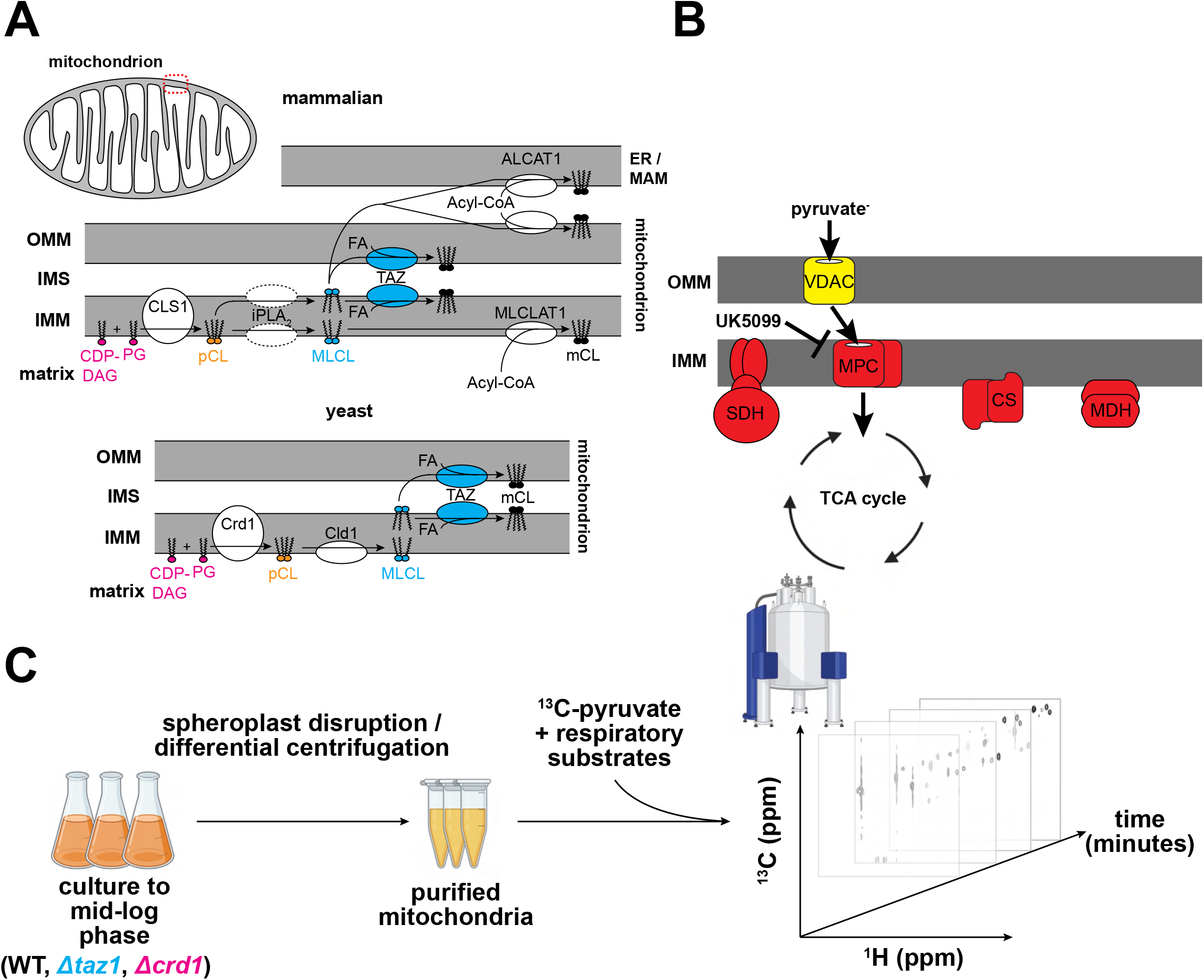
Cardiolipin (CL) remodeling and pyruvate uptake in mitochondria. *(A) Steps for CL biosynthesis and remodeling in mammals (upper) and yeast (lower)*. Mitochondria consist of an outer (OMM) and inner (IMM) membrane, which delineates the intermembrane space (IMS) from the matrix. Precursor CL (pCL) with variable acyl chain composition is synthesized *de novo* on the IMM by the joining of cytidine diphosphate diacylglycerol (CDP-DAG) to phosphatidylglycerol (PG) by cardiolipin synthase (CLS1/Crd1). CL then undergoes acyl chain remodeling first by the action of a deacylase (presumed to be a member of the iPLA_2_ family in mammals, Cld1 in yeast) to form monolyso-cardiolipin (MLCL). Then, a donor fatty acid (FA) is catalytically joined to MLCL by the transacylase tafazzin (TAZ) to form mature CL (mCL). Alternatively, in mammals, ALCAT1 can remodel mCL in the OMM or mitochondrial associated ER membrane (ER/ MAM) via utilization of a donor acyl tail from acyl-CoA. MLCLAT1 can also form mCL from acyl-CoA in the IMM. *(B) Mitochondrial pyruvate uptake and metabolism*. Cytosolic pyruvate is first taken up by mitochondria via the nonselective voltage-dependent anion carrier (VDAC) protein, and subsequently imported directly into the matrix by the selective mitochondrial pyruvate carrier (MPC). Pyruvate is then metabolized by enzymes of the TCA cycle, which includes the integral IMM succinate dehydrogenase (SDH) complex, as well as the peripherally-bound citrate synthase (CS) and malate dehydrogenase (MDH) proteins. The covalent inhibitor UK5099 blocks pyruvate uptake into the mitochondrial matrix by binding MPC. *(C) Schematic of in-organello real-time NMR approach to monitor mitochondrial metabolic flux*. First, mitochondria are isolated from WT, *Δtaz1*, and *Δcrd1* yeast strains by spheroplast disruption and differrential centrifugation. Then, mitochondria are incubated with ^13^C_3_ -pyruvate and respiratory substrates. Finally, a series of 2D ^1^H-^13^C HSQC NMR experiments are recorded over time to track the flux of the ^13^C-label.

Key metabolic enzymes are embedded in or are associated with the IMM (Fig. 1B). Although only one enzyme complex of the tricarboxylic acid (TCA) cycle is embedded in the IMM (succinate dehydrogenase, SDH) ^24^, many others are peripherally associated with the matrix-facing side of the IMM such as citrate synthase (CS) and malate dehydrogenase (MDH)^3^. These membrane-associated enzymes are believed to assemble into supra-molecular complexes known as metabolons that facilitate substrate channeling, where the product of one enzyme is passed directly to the next enzyme in the cycle ^25,26^. Another integral membrane protein is the mitochondrial pyruvate carrier (MPC) ^27^, a protein assembly necessary for pyruvate transport across the IMM. Therefore, we reasoned that the activities of these enzymes/complexes associated with the IMM could be perturbed by defects in CL biosynthesis since CL is a critical and abundant component of the IMM. Indeed, previous studies have demonstrated a link between changes in IMM lipid composition and changes in metabolite levels ^28^, particularly of metabolites involved in the TCA cycle ^29-32^. However, these past studies have focused on metabolites in plasma or whole-cell samples using primarily mass-spectrometry based methods, which typically only provide a single temporal snapshot of the metabolic state ^28-32^.

To obtain kinetic information of mitochondrial metabolomic flux, we have utilized an *in-organello* real-time Nuclear Magnetic Resonance (NMR) approach. NMR offers distinct advantages for metabolomics studies due to its non-invasive nature ^33,34^. In contrast to ‘one-shot’ mass spectrometry experiments where samples are destroyed, NMR makes it possible to monitor changes in metabolite levels as a function of time. Moreover, metabolites can be studied *in situ* without the need for solvent extraction that can potentially alter molecules and/or their concentrations. Finally, the use of ^13^C-labeling prevents interference from unlabeled molecules, thereby allowing unambiguous tracing of interconverting metabolites.

Here, we investigated the metabolism of ^13^C_3_-pyruvate, the central metabolite linking glycolysis and respiration, in purified mitochondria isolated from both WT yeast and isogenic strains defective in CL biosynthesis and remodeling (Fig. 1C). We used 2D ^1^H-^13^C heteronuclear single quantum coherence spectroscopy (HSQC) experiments to trace the ^13^C label and obtained additional 1D ^1^H nuclear Overhauser effect spectroscopy (NOESY) data to monitor the concentrations of nucleotides involved in energy metabolism. In our 2D ^1^H-^13^C HSQC experiments, we observed ^13^C-labeled metabolites that were distributed in the TCA cycle, and the synthesis of branched chain amino acids (BCAAs), fusel alcohols, and ergosterol. Additionally, the 1D ^1^H-NOESY experiments allowed us to monitor the flux of important energy nucleotides such as ATP, AMP, NAD+, and NADH. Finally, we compared how metabolite concentrations differed between mitochondria from WT yeast and *Δtaz1* and *Δcrd1* mutant strains lacking enzymes necessary for proper CL biosynthesis and remodeling, respectively. Taken together, these results offer new insights into how CL may modulate metabolism, and how an accumulation of MLCL (as in the case of BTHS) can be associated with defects in mitochondrial metabolism. Moreover, they also suggest that our real-time *in-organello* NMR approach is readily translatable and may be useful for studying mitochondrial metabolism in mammalian-based models of human disease.

## RESULTS

### Isolated mitochondria are active and show consistent purity among strains

Three *Saccharomyces cerevisiae* strains were used in this study, all isogenic to GA74-1A^35^. As confirmed in our previous studies, mitochondria from the WT strain have the normal complement of CL (∼10% of total phospholipids) with mostly unsaturated fatty acids; those from the Δ*crd1* strain have essentially no CL, but a buildup of its precursor, PG; and those from the Δ*taz1* strain have reduced CL content, an increased MLCL:CL ratio, and more saturated acyl chains ^35,36^. Thus, we isolated mitochondria from these three yeast strains across three independent isolation procedures per strain to ascertain the impact of IMM CL content on pyruvate metabolism.

To characterize our isolated mitochondria preparations, we first blotted our mitochondrial fractions against outer mitochondrial membrane (OMM) markers Tom40 and Tom70 to quantitatively assess their mitochondrial content, as the expression levels of OMM proteins is typically not affected by altered CL composition ^37^ (Fig. 2A, Fig. S1). Across our titrations, we observe similar levels of Tom70 and Tom40 in both our WT and *Δcrd1* mitochondria, whereas their abundance seemed to be slightly, but not significantly, decreased in the *Δtaz1* mitochondrial fractions (Fig. 2B).

**Figure 2:**
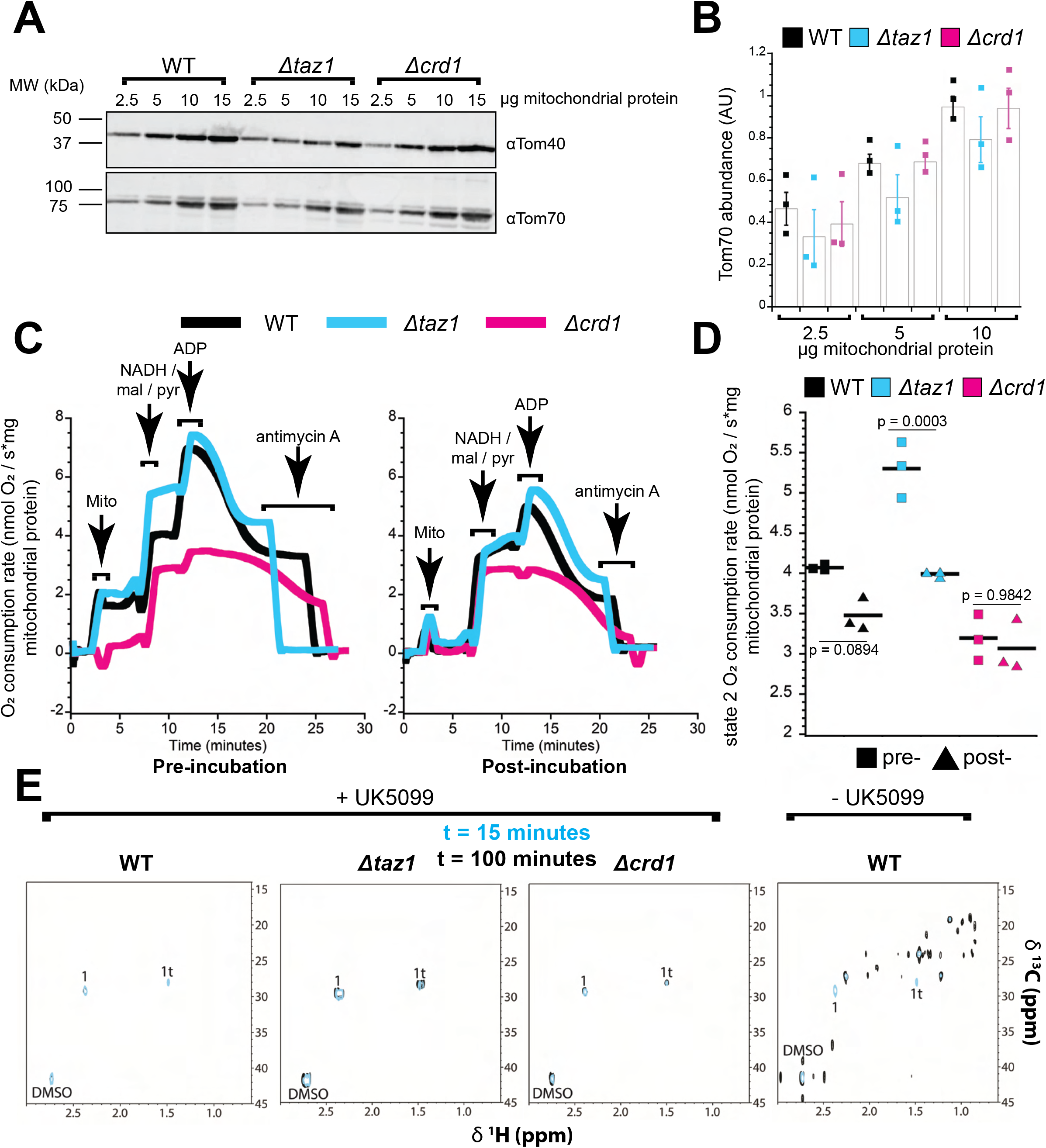
Purity and activity of mitochondrial isolated from yeast strains defective in CL biosynthesis. *(A) Representative Western blots against mitochondrial markers (Tom40 and Tom70)*. Uncropped blots for Tom70 are included in Supplementary Figure S1. *(B) Quantification of Tom70 abundance*. Tom70 band intensity was quantified in ImageJ from n = 3 independent replicates originating from two independent mitochondrial isolations for each strain. Values were normalized to signal obtained from loading 15 µg protein from WT mitochondria. *(C) Respirometry measurements prior to (pre-) and following (post-) incubation at 30°C for 65 minutes*. Shown here are representative traces. *(D) Quantification of state 2 respiration rate*. State 2 respiration pre-and post-65 minutes of incubation at 30*°*C was measured following addition of malate, pyruvate, and NADH for n = 3 independent biological replicates. p-values were obtained from one-way ANOVA and Tukey’s post-hoc test. *(E)* ^1^*H-*^*13*^*C3 HSQC spectra obtained following t=15 (cyan) and t=100 (black) minutes of incubation with* ^*13*^*C*_*3*_ *-pyru-vate*. Left panels were acquired in the presence of UK5099, which blocks mitochondrial pyruvate uptake through the MPC.

We next used high-resolution respirometry to analyze mitochondrial oxygen consumption activity in these strains as a metric of mitochondrial function (Fig. 2C, Fig. S2A). We chose to measure oxygen consumption rates pre-and post-65-minute incubations at 30°C to recapitulate the time window for performing the NMR metabolomic flux assays. Basal respiration rates (State 2 respiration induced by addition of pyruvate, malate, and NADH, Fig. 2D) did not significantly change for WT or *Δcrd1* across this time interval, whereas *Δtaz1* showed a slight yet significant decrease (p = 0.0003). Moreover, the basal oxygen consumption rates for *Δcrd1* and WT were similar, whereas the basal oxygen consumption rate was significantly elevated for *Δtaz1* (p = 5E-4 and 0.1671 for pre-and post-incubation measurements, respectively), likely due to increased uncoupling in the Δ*taz1* mutant^38^. Hence, although we observed some strain-specific effects on respiration due to prolonged 30°C incubation, all samples were respiration-competent.

During these time courses, we also measured State 3 and State 4 respiration rates (Fig. S2B). Relative to WT, *Δcrd1* demonstrated reduced state 3 and state 4 respiration rates for both pre-and post-incubation measurements (State 3: p = 1.5E-7, 4.7E-5; State 4: p = 1.3E-6,7.9E-5 for pre-and post-incubation measurements, respectively), whereas *Δtaz1* was similar except for increased pre-incubation state 4 respiration rate (p = 7.7E-4). Respiratory control ratios (RCRs, State 3/State 4) were similar between *Δtaz1* and WT for post-incubation measurements, but pre-incubation Δ*taz1* did display a decreased RCR compared to WT due to its increased State 4 rate. In contrast, the RCR for *Δcrd1* post-incubation was significantly elevated in comparison to WT, due to very low state 4 respiration likely caused from the stress imposed by forcing an ADP phosphorylation cycle after incubation. Following the 65-minute incubation, we observed a consistent decrease in respiration rate during both State 3 and State 4 respiration in all strains; consequently, this caused a significant increase in the RCR for all strains post-incubation.

Thus, we concluded that mitochondria from all strains maintained functional activity after an extended incubation at 30°C, and in line with previous reports ^35^, they displayed differences in their functional competency. Although prolonged 30°C incubation of mitochondria isolated from all three strains affected their ability to respond to phosphorylating conditions, they all remained capable of respiring and were impacted similarly. We therefore reasoned that they would retain reasonable functional integrity under the conditions required for our real-time NMR measurements.

Having verified that our mitochondria maintain activity over prolonged periods of time, we sought to characterize their metabolic flux using NMR. We collected 2D ^1^H-^13^C HSQC spectra of WT, *Δtaz1*, and *Δcrd1* mitochondria in the presence of 3 mM ^13^C_3_-pyruvate and respiratory substrates (see ‘Methods’). We first controlled for mitochondrial-specific pyruvate uptake and metabolism using the covalent MPC inhibitor UK5099 ^39^ (Fig. 2E). To this end, we collected spectra after t=15 and 100 minutes, where t=0 refers to the time of pyruvate addition, and the 15 minutes accounts for the NMR dead-time. At both timepoints in all three strains, we observed no change in the number of peaks in the presence of UK5099 (Fig. 2E, left), with all the peaks arising from either pyruvate or the UK5099 vehicle (DMSO). However, in the absence of UK5099 in WT mitochondria, we observed a significant increase in the number of cross peaks at the t=100 timepoint (Fig. 2E, right). The results verified that i) the IMM was intact in all the strains (as pyruvate can only enter the mitochondrial matrix through the MPC), ii) no contaminating cytosolic and/or ER components metabolized the pyruvate, iii) there was no interference from unlabeled cellular components, and iv) that several metabolites incorporating the ^13^C label could be observed after 100 minutes.

### Tracking the fate of mitochondrially metabolized ^13^C_3_-pyruvate

Given that we observed clear signs of pyruvate metabolism after t=100 minutes without an MPC inhibitor present, we then conducted a series of ^1^H-^13^C HSQC experiments throughout this time window. Figure 3 shows representative time points from a series of ^1^H-^13^C HSQC experiments used to track ^13^C_3_-pyruvate metabolism following mitochondrial uptake. In the earliest experiments the spectra are dominated by pyruvate (peak 1) and its enol tautomer (peak 1t) ^40^. Over the 90-minute course of the experiment, some 17 new peaks appear in the spectra that correspond to the 12 metabolites listed in Table 1. Assignments for the signals were based on metabolite and small compound NMR chemical shift databases ^41-47^, together with experiments in which mitochondria samples were matched to known standards, as described in the Methods section. Control ^13^C-HSQC experiments under the same conditions with unlabeled pyruvate ruled out that our set of assigned resonances could be from background 1% natural abundance ^13^C of unlabeled metabolites at large concentrations (data not shown).

**Table 1:** Mitochondrial metabolites derived from ^13^C3 -pyruvate detected in ^1^H-^13^C HSQC experiments^a,b^.

| Metabolite | NMR Peak | Pathway | Ref. | WT Range (fmol/ng)<sup>b</sup> | $\Delta$ time | $\Delta taz1$ | $\Delta crd1$ |
| --- | --- | --- | --- | --- | --- | --- | --- |
| Pyruvate/Tautomer | 1, 1t | Substrate |  | 295 to 0 | ↓↓ | 0 | 0 |
| Citrate | 12 | TCA | 62,102 | 1.6 to 0.3 | 0 | 0 | 0 |
| α-ketoglutarate | 14 | TCA | 62,102 | 2.7 to 0.5 | ↑↓ | ↓↑ | ↓0 |
| Succinate | 18 | TCA | 62,102 | 1 to 7.7 | ↑ | ↓ | ↓ |
| Glutamate | 10 | GDH & GOGAT | 68-70 | 0.13 to 0.5 | 0 | 0 | 0 |
| Acetolactate | **16, 17** | BCAA | 77,103 | 92 to 200 | ↑ | ↓ | ↓↓ |
| α-ketoisovalerate | 13, **15** | BCAA/Val | 77,103 | 10 to 43 | ↑ | ↓↓ | ↓↓ |
| α-isopropylmalate | **4, 7, 11** | BCAA/Leu | 77,103 | 0.94 to 1.7 | ↑ | ↓ | ↓ |
| Valine | **6, 8** | BCAA/Val | 77,103 | 0.12 to 0.6 | ↑↑ | 0 | ↓ |
| Acetoin | 5 | Butanediol | 52,76 | 0 to 1.6 | 0 | ↑ | 0 |
| Mevalonate | 2 | Sterol | 51,81 | 0.9 to 1.25 | 0 | ↑↑ | 0 |
| Citramalate | 3 | Citramalate/Ile | 80,104 | 13 to 32 | ↑ | ↑ | 0 |
| Acetate | 9 | Pyruvate Decarboxylation | 40,53 | 3.7 to 7.8 | 0 | 0 | 0 |

<sup>a</sup>Columns from left to right: compound, NMR peak number, associated pathway, literature references for the pathway, concentration range over the course of the ~90 minute experiments, direction of change over time for WT, change in $\Delta taz1$ compared to WT, change in $\Delta crd1$ compared to WT. Double arrow indicates more than a 10-fold change in metabolite concentration, and a single arrow is less than a 10-fold change. For the mutants, 0 indicates no significant change, ↓ indicates a decrease, and ↑ indicates an increase in the concentrations of the given metabolite compared to WT. For metabolites with multiple NMR peaks, the average of the the peaks highlighted in bold were used for quantification. To save space, the kinetic profile for acetate is not shown in Fig. 5.
<sup>b</sup>The starting concentrations given are those calculated from the first NMR spectra, recorded after a ~15 minute deadtime from the start of the reaction. T=0 corresponds to when the 3 mM $^{13}\text{C}_3$ -pyruvate was added to the mitochondria.

**Figure 3:**
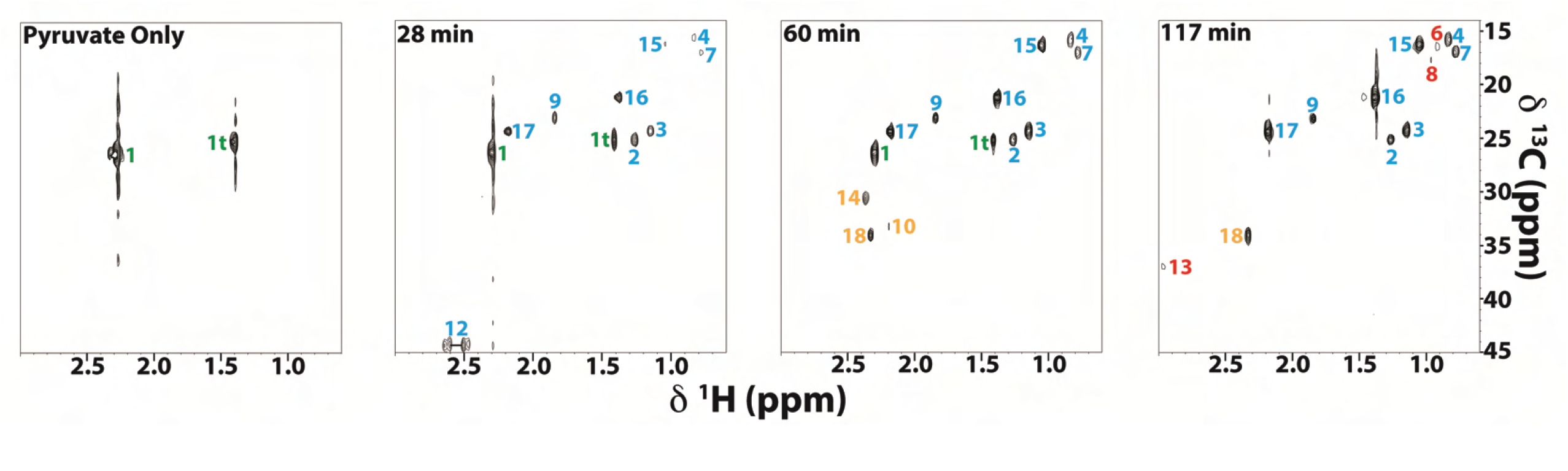
Representative ^13^C3 -pyruvate flux experiment in WT mitochondria. Each panel represents an individual 2D ^1^H-^13^C HSQC spectrum collected at the specified time over the course of the experiment. Numbers listed next to each peak correspond to NMR signals from the metabolites listed in Table 1, and are shown in different colors depending on the time of their first appearance in the spectra (green, initial; cyan, 28 min; orange, 60 min; red, 117 min). “1t” corresponds to the enol tautomer of pyruvate.

The metabolites detected correspond to pathways in yeast mitochondria for which pyruvate or pyruvate-derived TCA cycle intermediates serve as the entry point (Fig. 4). Within the TCA cycle, we detected the three metabolites citrate, αKG, and succinate. Some TCA metabolites may not have been observed because they are either too short-lived, bound to proteins too large for NMR detection, or because they occur outside the spectral window sampled in the NMR spectra (e.g. if the ^13^C-label is incorporated in carbonyl carbons). In addition to TCA cycle metabolite intermediates, we also saw metabolites from branching cataplerotic pathways emanating from the TCA cycle hub. For example, glutamate is produced through transamination of the TCA cycle metabolite αKG^48^. The metabolites acetolactate, α-ketoisovalerate, α-isopropylmalate and valine, all of which were observed in our HSQC experiments, are in the BCAA synthesis pathways. Unlike yeast, humans are incapable of synthesizing BCAAs and these essential amino acids must thus be supplemented from the diet ^49^. Other metabolites we observed include: citramalate, which is a precursor in isoleucine biosynthesis (formed by pyruvate and acetyl-CoA ^50^), mevalonate, which is an intermediate in the yeast sterol pathway leading to ergosterol ^51^, acetoin, which can be produced from acetolactate through the yeast ‘fusel’ alcohol pathway under stress conditions ^52^, and acetate, which can form from pyruvate decarboxylation as a result of the highly-oxidizing environment of mitochondria ^53^. All the metabolites described above have been documented to occur in yeast mitochondria.

**Figure 4:**
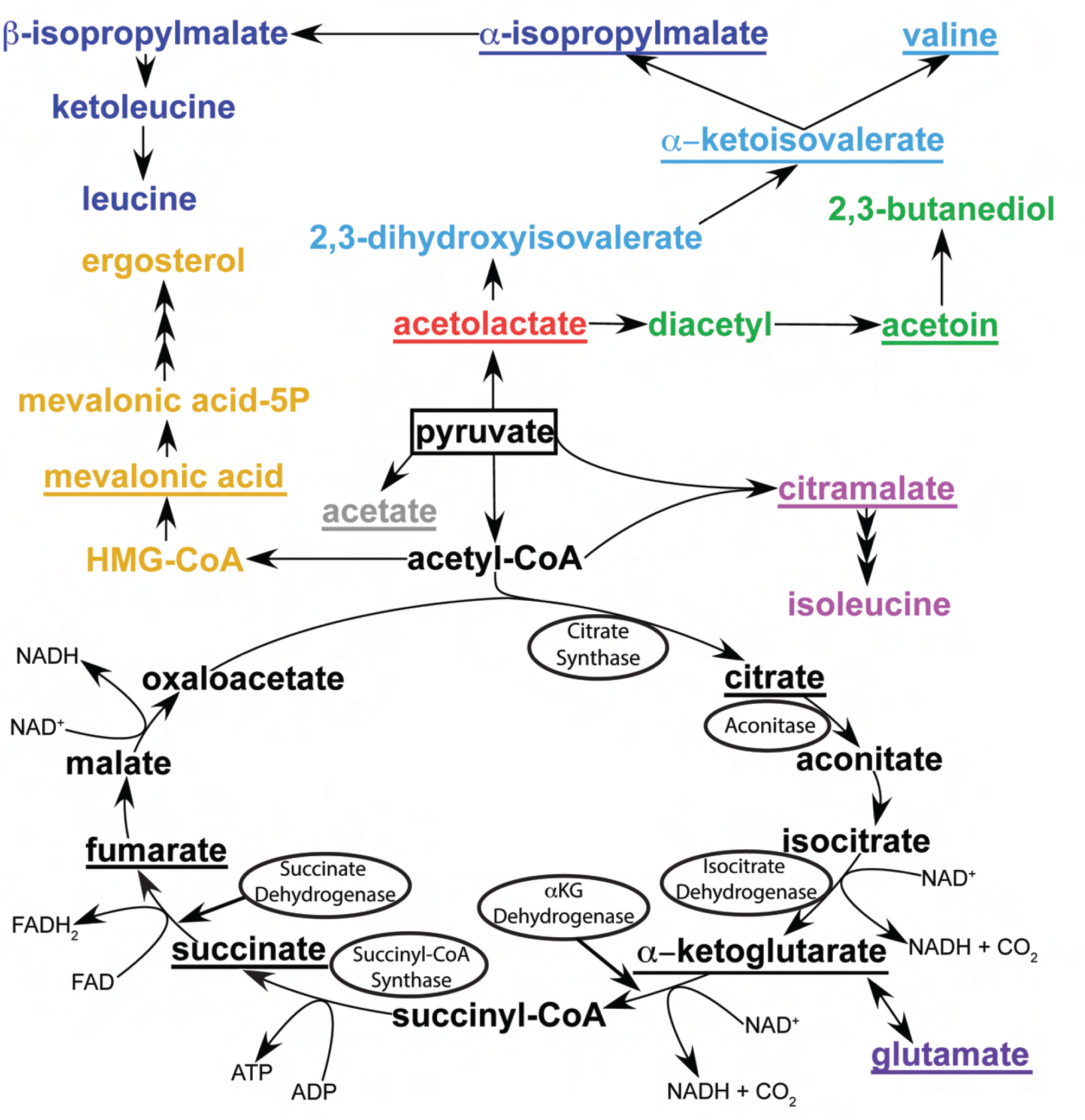
Pathways of pyruvate metabolism in yeast mitochondria. Metabolites derived from pyruvate that are observed in the NMR experiments are underlined. The starting substrate (pyruvate) is boxed. Tricarboxylic acid (TCA) cycle enzymes that are known to require CL for activity are circled. Color legend: black, TCA cycle; green, butanediol pathway; light/dark blue, valine and leucine arms of the branched chain amino acid (BCAA) pathway, respectively; pink, citramalate pathway leading to isoleucine; violet, glutamine oxoglutarate aminotransferase (GOGAT)/gluta-mine dehydrogenase (GDH) pathway; yellow, sterol pathway; grey, acetate formation through pyruvate decarboxylation (PD). Refer to Table 1 for references.

### Comparison of mitochondrial metabolic flux between WT and mutants with defective CL biosynthesis and remodeling

To investigate differences in metabolic flux between WT and the CL biogenesis and remodeling mutants Δ*taz1* and Δ*crd1*, we recorded ^1^H-^13^C HSQC data on isolated mitochondria from each strain following addition of ^13^C_3_-pyruvate (Fig. 5). Experimental dead-times between preparing samples and recording the first NMR spectra were on the order of ∼15 minutes. Spectra were sampled every ∼5.5 minutes for a total experiment time of ∼1.5 hours, as our controls (Fig. 2B) indicated the mitochondria remained active for at least 90 minutes.

**Figure 5:**
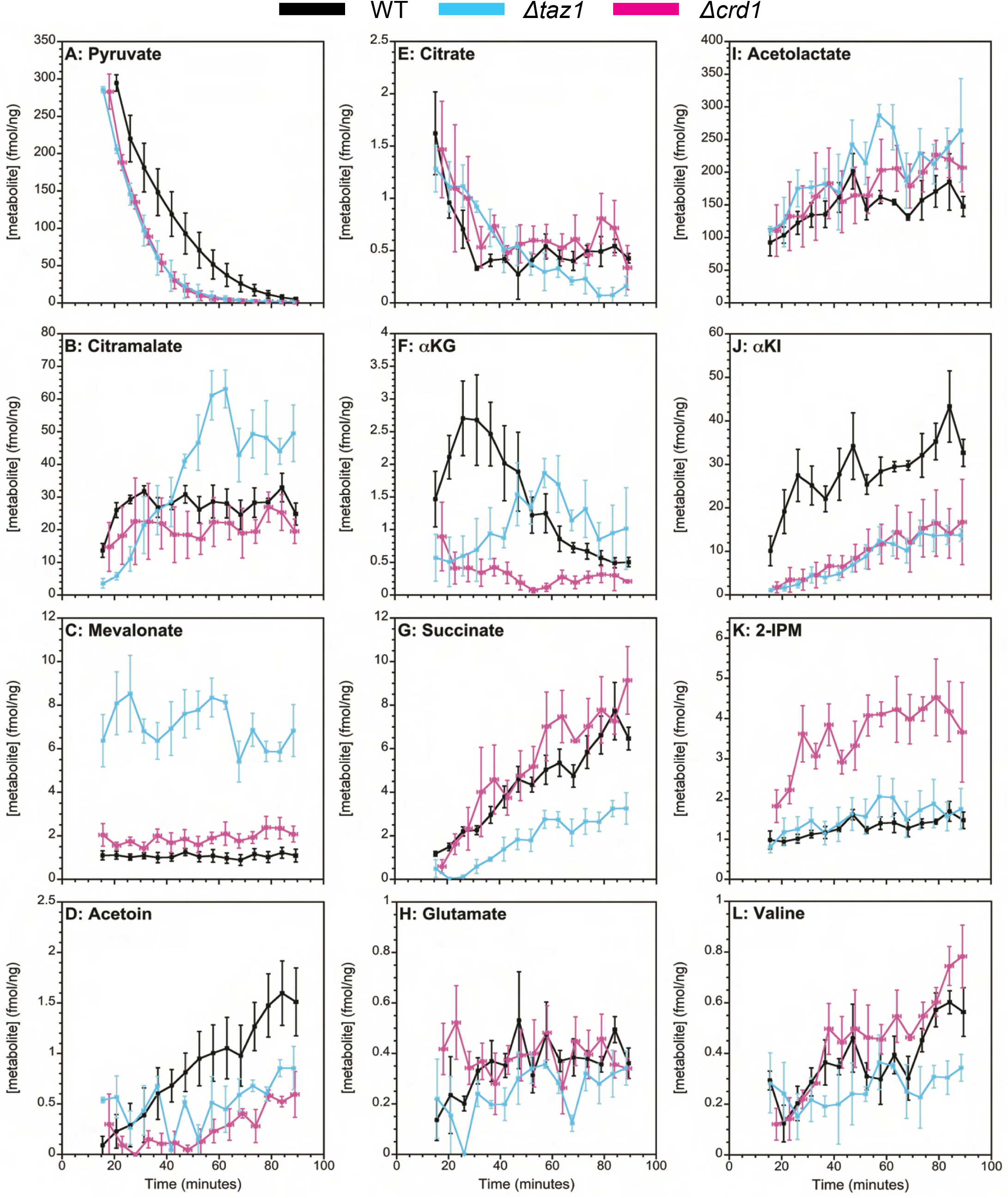
Kinetic traces of mitochondrial metabolite flux observed by 2D ^1^H-^13^C HSQC experiments. Data points are colored according to yeast strain: black, WT; cyan, *Δtaz1*; pink, *Δcrd1*. Y values (fmol metabolite per ng of mitochondrial protein) represent means ± standard error (SEM) from n ≥ 3 individual experiments. X-error bars represent variation (± SEM) in timing of the n ≥ 3 experiments. T=0 corresponds to the time of ^13^C_3_ -pyruvate addition. Panels are arranged so that (A-D) correspond to pathways for which a single metabolite is observed (A: pyruvate, B: citramalate, C: mevalonate, D: acetoin), (E-H) correspond to the TCA cycle and glutamate pathways (E: citrate, F: αKG, G: succinate, H: glutamate), and (I-L) corresponds to the BCAA pathways (I: acetolactate, J: αKI, K: 2-IPM, L: valine). For metabolites with multiple peaks (Table 1), the peaks used to determine concentrations were 1, pyruvate; an average of 16 and 17, acetolactate; 15, αKI; an average of 4 and 7, 2-IPM; and an average of 6 and 8, valine.

To initiate the reactions, a 3 mM concentration of the ^13^C_3_-pyruvate substrate was added to a sample that contained mitochondria corresponding to 1 mg of mitochondrial protein. Therefore, the calculated starting concentration of the pyruvate was 3,000 fmol/ng mitochondria. After the 15-minute dead-time of the experiment, the earliest pyruvate concentrations detected by NMR were around ∼300 fmol/ng, about 10-fold less than the theoretical starting concentration. Similar 3 mM substrate concentrations and starting levels for pyruvate were observed during comparable NMR experiments for mitochondria isolated from HCT116 human colon cancer cell^54^. Either most of the ^13^C_3_-pyruvate is metabolized in the 15-minute dead-time of the experiment before NMR data is collected, or the ^13^C label is incorporated into macromolecules and complexes too large to be seen by NMR (>∼30 KDa). Because of the size-filtering properties of NMR ^55,56^ our experiments only detect the pool of metabolites freely circulating in solution, whereas the fraction bound as cofactors in macromolecular complexes can be larger ^57,58^.

For the metabolites we observe, it is useful to distinguish between four types of flux profiles. (i) A decaying profile (for all newly detected metabolites except the ^13^C_3_-pyruvate substrate) implies that there must be an initial build-up of the compound that is not observed during the dead-time of the NMR experiments ^59^. These types of profiles are rare in our dataset, but an example occurs for citrate (Fig. 5E). A similar profile for citrate was also observed with mitochondria isolated from HCT116 cells ^59^. (ii) A second profile type displays an initial buildup phase followed by a decay which is demonstrated by WT with the TCA cycle intermediate α-ketoglutarate (αKG) (Fig. 5F). This suggests the αKG is metabolized more slowly than citrate, allowing it to accumulate before it eventually decays. (3) A flat profile, meanwhile, suggests a balance between the production and consumption of a metabolite, yielding an apparent steady state during the time course of the experiment. Such a profile is manifested for mevalonate (Fig. 5C).(4) Finally, succinate shows a profile with an apparent steady rise to an eventual plateau (Fig. 5G). This type of profile is also seen for succinate in HCT116 mitochondria ^59^, and suggests an increase in concentration during the time course of the experiment.

Metabolites involved in cyclical pathways such as the TCA cycle tended to show flat or decreasing profiles, whereas those at the ends of pathways tended to show profiles that rise to a plateau. For example, 2-isopropylmalate (2IPM) (Fig. 5K) is an intermediate in the synthesis of the amino acid leucine, but the biosynthetic steps following this compound occur in the cytosol ^60^; thus, this may account for the rise to a steady-state plateau during the course of our experiments in isolated mitochondria.

It is informative to compare the levels of the different metabolites shown in Figure 5. Excluding the starting substrate pyruvate, by far the largest concentrations of ∼200 fmol/ng occur for acetolactate (Fig. 5I), which is an entry point for both the BCAA and fusel alcohol pathways (Fig. 4). For comparison, the largest concentrations observed for any of the TCA pathway metabolites (Fig. 5E-G) are ∼8 fmol/ng for succinate. This suggests that under the conditions of our experiments, about 100-fold more pyruvate is used for BCAA synthesis than for the TCA cycle. Note that in these experiments the mitochondria are approaching state 4 respiration which would put lower demands on NADH synthesis, diverting carbon flux towards alternative metabolic pathways. In contrast, under uncoupled or state 3 respiration conditions, TCA flux may be more dominant ^61^. It is also possible that much more of the ^13^C label is residing in TCA cycle metabolites than we can visualize. Due to the potential lack of bottlenecks in a self-contained cyclic pathway with substrate channeling, the intermediate metabolites could be kept bound to large complexes and therefore be invisible to NMR detection. In contrast, the BCAA pathways which have endpoints that require non-mitochondrial enzymes could lead to metabolite build-ups. Finally, the measured concentrations of ^13^C-label derived from ^13^C_3_-pyruvate decreased the further the metabolite in a pathway was separated from the initial substrate. Thus, it is important to note that this may be a manifestation of ^13^C-label dilution with ^12^C, as the labeled carbons from the original substrate participate in an increasing number of metabolic reactions and are also dissipated in the form of CO_2_.

### ^1^H-NMR assessment of ATP and NADH energy metabolism

While the ^1^H-^13^C HSQC experiments in Figures 3,5 make it possible to follow the fate of the ^13^C-label derived from pyruvate, they do not provide information on the subset of important nucleotide metabolites involved in mitochondrial respiration and energy production such as ATP, AMP, NAD^+^ and NADH. We therefore investigated changes in the concentrations of these compounds under the conditions of our ^13^C-labeled studies by keeping all experimental variables identical, except substituting unlabeled ^12^C_3_-pyruvate for ^13^C_3_-pyruvate. NMR resonances associated with the nucleotides and other metabolites in the downfield region of the ^1^H NMR spectrum (Fig. 6) was monitored as a function of time following the addition of unlabeled ^12^C_3_-pyruvate. Note that these experiments detect both ATP/ADP and NAD^+^/NADH that were endogenous to mitochondrial samples, and those that were added exogenously in the reactions. Namely, the isolated mitochondria samples were supplemented with a respiratory substrate mix consisting of 0.25 mM malate, 0.25 mM NADH,0.1 mM ADP, 0.1 mM ATP in addition to 3 mM pyruvate. Moreover, we are only likely to observe free nucleotides, as nucleotides bound to large proteins or complexes would not be visible by NMR. For example, it has been previously reported that ∼80% of the cellular NAD^+^/NADH pool is immobilized and bound to proteins ^57^. Despite these limitations, we can still compare the mitochondrial metabolic turnover of nucleotides among the strains as identical concentrations of exogeneous nucleotides were added to each (Fig. 7).

**Figure 6:**
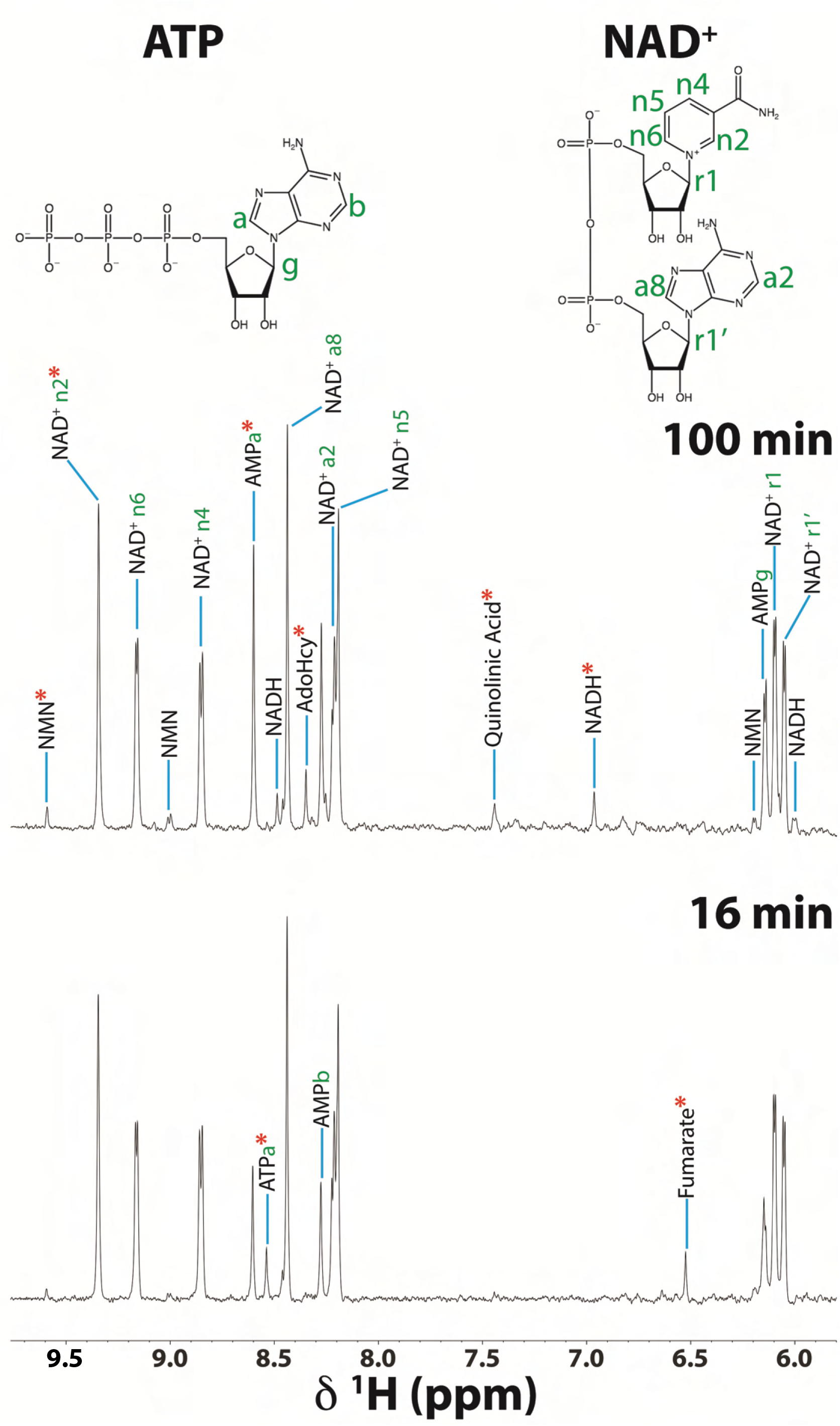
Downfield region of the 1D ^1^H NMR spectrum following pyruvate addition. Representative 1D ^1^H NOESY spectra of WT mitochondria at the start (bottom) and end (top) of the NMR timecourses using 3 mM unlabeled ^12^C_3_ -pyruvate in place of 3 mM ^13^C_3_ -pyruvate. Peaks are labeled with their indicated NMR assignments ^105^. Only peaks not appearing in the spectra at 100 min are labeled in the spectra at 16 min. The positions of protons corresponding to the NMR assignments of ATP and NAD+ are labeled in their structures. The red ^*^ symbols indicate NMR signals that were sufficiently isolated in the spectra to allow quantification of the respective metabolite concentration (see Fig. 7).

**Figure 7:**
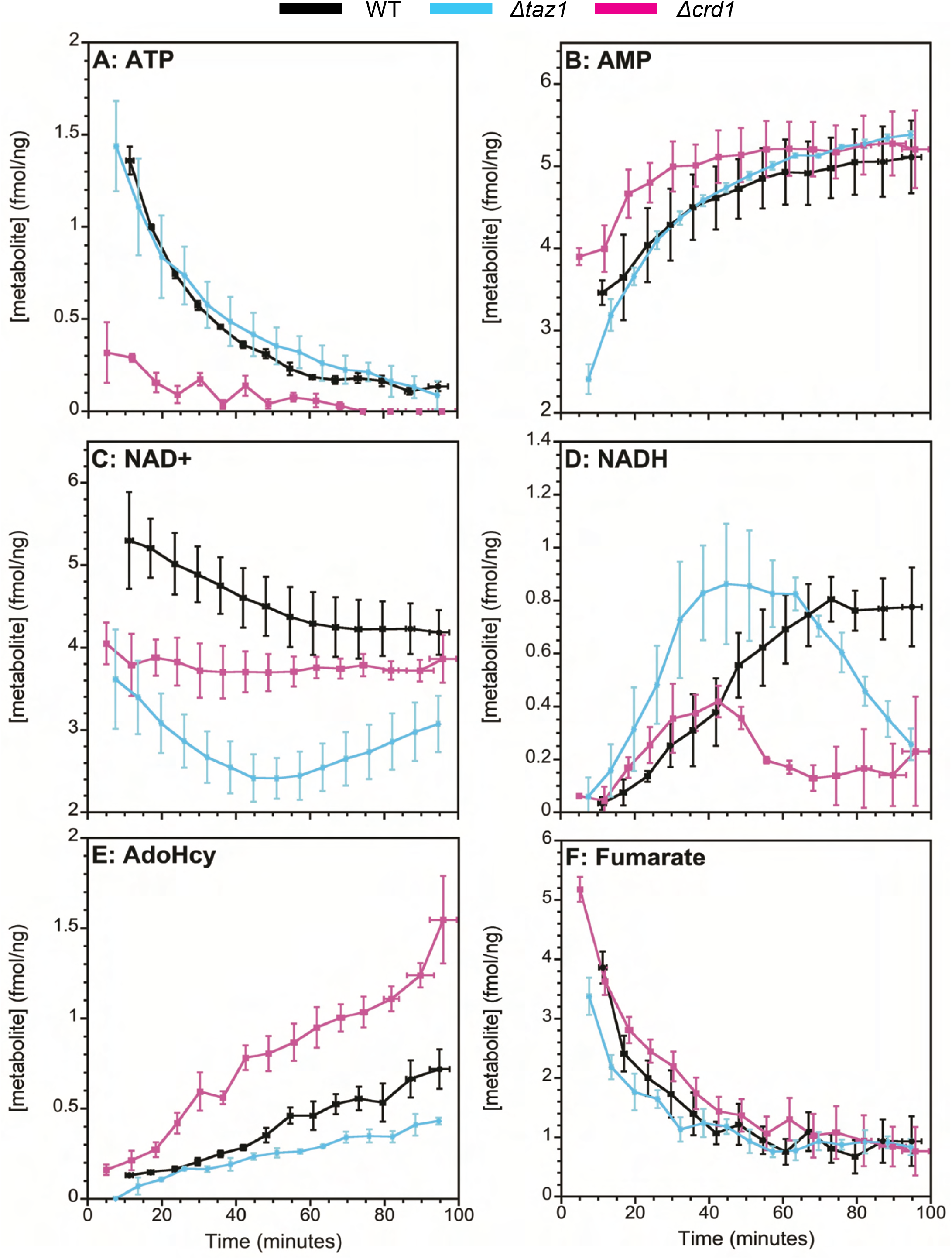
Mitochondrial metabolites observed by 1D ^1^H NOESY experiments. Kinetic profiles of selected metabolites obtained from the 1D ^1^H NMR data illustrated in Fig. 6. (A) ATP, (B) AMP, (C) NAD+, (D) NADH, (E) S-adenosyl homocysteine (AdoHcy), (F) fumarate.

In each turn of the TCA cycle, six electrons (three hydride ions) are transferred to three NAD^+^ molecules to form NADH (an additional pair of electrons in two hydrogen atoms are transferred to FAD to make FADH_2_). NADH is subsequently oxidized back to NAD^+^ (and FADH_2_ is oxidized back to FAD) as the reducing equivalents are donated to the electron transport chain (ETC) via reduction of Coenzyme Q10. Then, electrons originating from reduced Coenzyme Q10 are transferred to Complex III, cytochrome c, and Complex IV along the ETC to facilitate the asymmetric pumping of protons across the IMM; in turn, this generated gradient is used to synthesize ATP ^62^.

The levels of ATP produced through oxidative phosphorylation coupled to NADH oxidation are similar for WT and *Δtaz1* but are decreased about 6-fold at the start of the experiment for *Δcrd1* (Fig. 7A). Furthermore, AMP levels show a more rapid initial increase for *Δcrd1* compared to WT and *Δtaz1* immediately following the NMR dead-time, but all strains eventually plateau to similar concentrations (Fig. 7B). The fact that net NAD^+^ levels decrease (Fig. 7C) and net NADH levels increase (Fig. 7D) over the course of our experiments indicates that TCA cycle activity is causing the matrix to shift to a more reducing environment. However, we observed that this transition to a more reducing environment is slower in Δ*crd1* mitochondria than WT mitochondria. With Δ*taz1*, the levels of NAD^+^ instead fall and then increase after 60 minutes. This pattern is further reflected in the NADH levels that increase up to 60 minutes and then decrease.

We were also able to detect two additional metabolites in the 1D ^1^H NMR experiments, S-adenosyl homocysteine (AdoHcy) and fumarate. AdoHcy is produced when S-adenosyl-methionine (AdoMet) acts as a methyl donor in several lipid synthesis pathways ^63,64^. Levels of AdoHcy increased over time in our experiments, with the Δ*crd1* mutant showing a∼3-fold larger increase than WT or Δ*taz1* (Fig. 7E). The blocking of CL biosynthesis in the Δ*crd1* strain is likely causing a buildup of its precursor cytidine diphosphate diacylglycerol (CDP-DAG), which could lead to an increase in PC synthesis, and thus an increase in AdoHcy levels ^65^. Finally, we detected the TCA cycle intermediate fumarate from its well-resolved ^1^H shift at 6.5 ppm. This compound was not observed in our ^13^C-HSQC spectra because the symmetric methine CH carbons of fumarate resonate at 138 ppm, outside of our ^13^C spectral window running between -3 and 76 ppm. Fumarate levels show similar flux profiles between WT and the CL remodeling mutants. A summary of the metabolites detected in our 1D experiments can be found in Table 2.

**Table 2:**
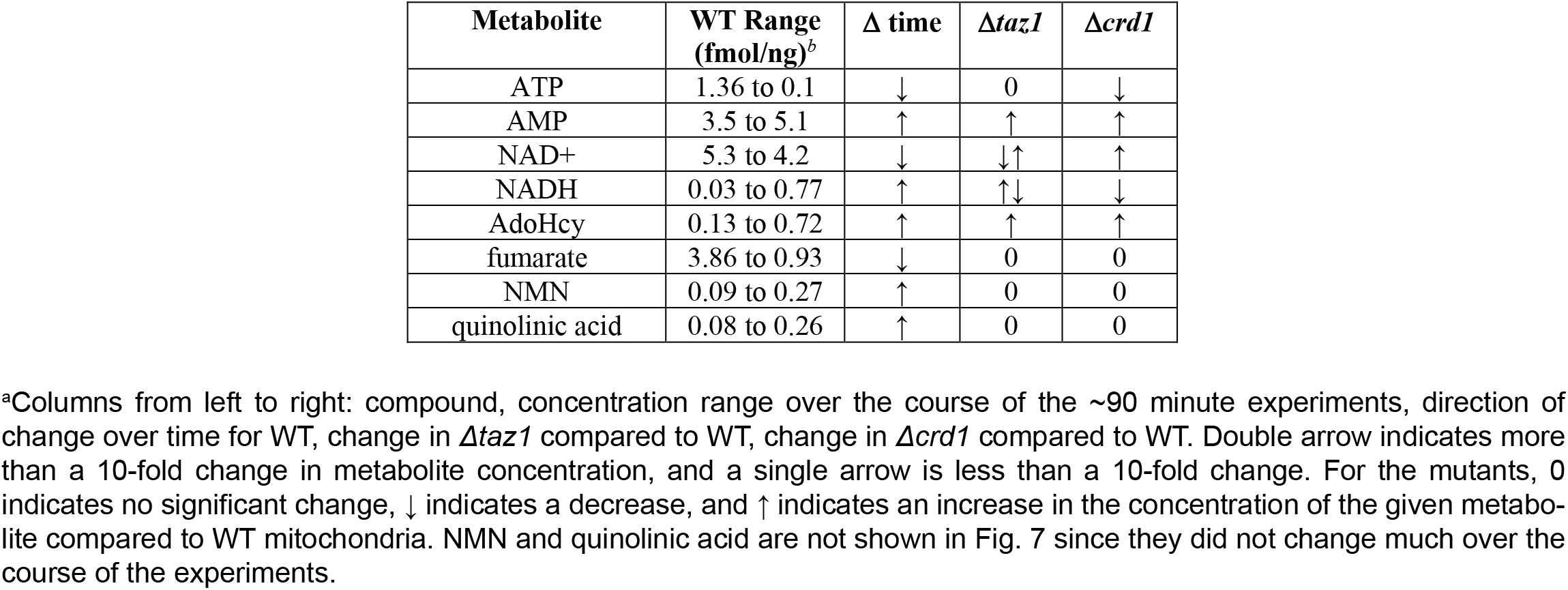
Mitochondrial metabolites detected in 1D ^1^H NOESY experiments^a^.

| Metabolite | WT Range (fmol/ng)<sup>b</sup> | Δ time | Δ*taz1* | Δ*crd1* |
| --- | --- | --- | --- | --- |
| ATP | 1.36 to 0.1 | ↓ | 0 | ↓ |
| AMP | 3.5 to 5.1 | ↑ | ↑ | ↑ |
| NAD<sup>+</sup> | 5.3 to 4.2 | ↓ | ↓↑ | ↑ |
| NADH | 0.03 to 0.77 | ↑ | ↑↓ | ↓ |
| AdoHcy | 0.13 to 0.72 | ↑ | ↑ | ↑ |
| fumarate | 3.86 to 0.93 | ↓ | 0 | 0 |
| NMN | 0.09 to 0.27 | ↑ | 0 | 0 |
| quinolinic acid | 0.08 to 0.26 | ↑ | 0 | 0 |

<sup>a</sup>Columns from left to right: compound, concentration range over the course of the ~90 minute experiments, direction of change over time for WT, change in Δ*taz1* compared to WT, change in Δ*crd1* compared to WT. Double arrow indicates more than a 10-fold change in metabolite concentration, and a single arrow is less than a 10-fold change. For the mutants, 0 indicates no significant change, ↓ indicates a decrease, and ↑ indicates an increase in the concentration of the given metabolite compared to WT mitochondria. NMN and quinolinic acid are not shown in Fig. 7 since they did not change much over the course of the experiments.

## DISCUSSION

### Roles of the compounds derived from ^13^C_3_-pyruvate in yeast mitochondrial metabolism

NMR-based *in-organello* studies of ^13^C_3_-pyruvate metabolic flux were first done with mitochondria isolated from the human colon cancer cell line HCT116 ^54^. In this previous study, citrate and succinate were the only two TCA cycle metabolites identified for the HCT116 mitochondria. The flux kinetic profiles of these two metabolites were similar to those we observe in yeast, with citrate decreasing and succinate increasing with time. Concentrations of succinate levels were comparable in a range of 1 to 6 fmol/ng for the two sources of mitochondria, but citrate levels were ∼30-fold higher in human cancer cell mitochondria (30 to 15 fmol/ng) than in yeast (Table 1). Citrate levels are expected to be lower in cancer cells ^59^, so the low citrate levels we observed in yeast could be due to species-related differences. We identified the additional TCA cycle metabolite αKG (Table 1, Fig. 3), which we believe was incorrectly assigned in the HCT116 mitochondria study as glutamine, since glutamine is only synthesized in the cytosol of most organisms (except uricotelic vertebrates, where it is used to detoxify mitochondrial ammonia ^66,67^). Finally, we observed the TCA metabolite fumarate in 1D ^1^H NMR experiments (Fig. 7), which decreased from an initial level of 3.9 to 0.9 fmol/ng over the time course of the experiments. An important difference between the HCT116 study and our yeast study is that acetyl phosphate was detected in the HCT116 mitochondria as an oxidation product of ^13^C_3_-pyruvate, but not in mitochondria with a knockout of the p53 gene ^54^. We, however, did not observe acetyl phosphate in yeast mitochondria. Thus, either the compound is absent in yeast mitochondria and/or too short-lived to be observed. The TCA cycle metabolites cis-aconitate, isocitrate, succinyl-CoA, malate, and oxaloacetate were also not detectable by NMR in both yeast and HCT116 mitochondria.Outside the TCA cycle, the metabolites acetate and glutamate were seen with both HCT116 ^59^ and yeast mitochondria (Table 1, Fig. 3). Acetate and glutamate concentrations were∼30-fold higher in the HCT116 mitochondria ^59^ than in yeast (Table 1). Acetate can be produced through oxidative decarboxylation of pyruvate by hydrogen peroxide ^53^, or alternatively as an enzymatic byproduct of pyruvate dehydrogenase – the enzyme that forms acetyl-CoA from pyruvate ^40^. Glutamate in yeast can be synthesized by mitochondrial enzymes such as glutamic dehydrogenases GDH2 ^68^ or GDH3 ^69^ and by the glutamate synthetase (GOGAT) GLT1^70^. These enzymes use the TCA cycle intermediate αKG as a precursor for transamination, with glutamine imported from the cytosol acting as the sole nitrogen source ^48,71-74^.

Of the 12 metabolites monitored in this work, seven compounds are in pathways unique to yeast. The strongest NMR signals in our ^1^H-^13^C HSQC spectra besides ^13^C_3_-pyruvate are from acetolactate. The mitochondrial enzyme acetolactate synthase condenses two molecules of pyruvate to give 2-acetolactate and CO_2_ ^75^. Acetolactate is the entry point for the yeast mitochondrial synthesis of the BCAA valine ^76^. In the mitochondrial valine synthesis pathway, we observe the additional metabolites α-ketoisovalerate and valine (Table 1, Fig. 4). The compound α-isopropylmalate is an intermediate in the synthesis of the BCAA leucine. The metabolite is synthesized in mitochondria by the long isoform of α-isopropylmalate synthase Leu4 ^77^, but it is subsequently transported to the cytosol via the mitochondrial oxaloacetate carrier Oac1p ^60^, where most of the leucine synthesis pathway occurs ^60,76,77^.

The last three metabolites unique to yeast are acetoin, citramalate, and mevalonate (Table 1). Acetoin can be produced from acetolactate in the Ehrlich ‘fusel alcohol’ pathway ^76,78^ by the butanediol dehydrogenase BDH1 ^79^. However, the Ehrlich pathway that produces higher alcohols from assimilated amino acids typically occurs in the cytosol ^76^. Despite this, a recent report suggests that mitochondrial compartments could act as fusel-alcohol generators by catabolizing BCAAs under conditions of elevated amino acid stress ^52^. Alternatively, acetoin could be present in mitochondria as a side-product of pyruvate dehydrogenase since it occurs at small levels of ∼1 fmol/ng, and since literature NMR spectra show that acetate, acetolactate, and acetoin can be produced during pyruvate turnover by the enzyme ^40^. Citramalate is formed from pyruvate and acetyl-CoA and may be involved in yeast mitochondrial isoleucine synthesis ^50,80^. Mevalonate is produced from pyruvate via acetyl-CoA in the mitochondrial sterol pathway ^81^. The sterol pathway eventually produces ergosterol ^51^, the yeast analog of cholesterol.

Thus, all the metabolites we identified by NMR can be traced to yeast mitochondrial metabolism originating from our starting substrate pyruvate.

### Comparison of TCA metabolites in WT and CL biosynthesis and remodeling mutant mitochondria

Pyruvate consumption was similar between WT and CL biosynthesis and remodeling deficient mitochondria, with exponential decay constants equal to ∼16 minutes (Fig. 5A). Although pyruvate visually appears to have been metabolized faster in Δ*taz1* and Δ*crd1* mitochondria relative to WT, the differences were not statistically significant (p ≥ 0.05). The similar pyruvate turnover profiles thus suggest that neither pyruvate transport through the MPC, nor the initial steps in pyruvate catabolism, are affected by CL. In the TCA cycle, the kinetic profiles of the earliest compound in the cycle, citrate (Fig. 5), and the last compound, fumarate (Fig. 7), were also similar between WT and the mutants. However, it is important to note that fumarate was observed only in the 1D ^1^H NMR experiments, and as such may not be directly comparable to the three TCA cycle metabolites that are detected through the ^13^C-label (citrate,αKG, and succinate). Despite this, the measured concentrations of fumarate agreed with the other TCA cycle metabolites.

Of the three, αKG in WT and Δ*taz1* showed a biphasic profile characterized by an initial rise followed by a gradual decrease, with the peak αKG concentration lagging by about 20 minutes in Δ*taz1* compared to WT. The αKG concentrations for Δ*crd1* were relatively constant and 3-fold lower than the peak levels for WT or Δ*taz1*. These differences may be due to perturbation of the SDH complex, the only integral IMM TCA cycle enzyme that participates directly in the electron transport chain ^82^. αKG can be converted to glutamate by the glutamate dehydrogenase and glutamate synthase enzymes ^68-70^. The levels of glutamate were low and equivalent among the three strains, thus indicating that these pathways are unlikely to be the source of the differences in αKG profiles. The next observed metabolite in the TCA cycle following αKG, succinate (Fig. 5G) showed lowered levels in Δ*taz1*, which may be related to the lag observed with αKG. Lowered levels for TCA cycle metabolites in Δ*taz1* mitochondria may also be related to higher levels of cataplerotic metabolites such as citramalate and mevalonate (Fig. 5B and Fig. 5C, respectively). In contrast, the succinate levels for the Δ*crd1* mitochondria were comparable to WT, even though the mutant showed much lower levels of the precursor αKG. Therefore, the absence of CL (Δ*crd1*) or its abrogated remodeling (Δ*taz1*) had different effects on the enzymes that regulate TCA metabolite flux. Of particular interest, the TCA metabolites αKG, succinate, and fumarate are increasingly recognized for their roles in regulating the expression of genes that determine cell fate, although this role may not directly pertain to yeast ^62^.

### Effects of deficient CL biosynthesis and remodeling on nucleotide energy metabolites

We observed differences among the three strains for adenine and pyridine nucleotide concentrations and kinetics (Fig. 7, Table 2). ATP levels were similar in WT and Δ*taz1* but much lower in Δ*crd1*. The measured ATP/AMP ratios in our yeast mitochondria were significantly lower than previous results obtained in mammalian mitochondria ^83,84^. One possible explanation for this discrepancy could be the presence of adenylate kinase activity in our mitochondrial samples ^85^; alternatively, our NMR-based approach may be insensitive to nucleotides bound in large complexes as noted previously. Lastly, different yeast growth conditions and/or isolation protocols could also contribute to different mitochondrial metabolic activities and levels of endogenous metabolites. The individual concentrations for AMP and ATP, however, were comparable to values found in the literature ^86-88^, with yeast mitochondrial ATP concentrations of ∼1.5-2 fmol/ng being reported for WT mitochondria ^86^.

Differences in NAD^+^ and NADH levels suggested impaired energy metabolism and/or redox regulation in the CL biosynthesis and remodeling mutants compared to WT. Mitochondrial NAD+/NADH ratios of ∼5-10 have been previously reported, and these values agreed with our measurements ^89,90^. Since NADH is produced by the TCA cycle, our data suggest that this pathway may be impaired in Δ*taz1* and Δ*crd1* mitochondria, which is in line with a previous report ^32^. These differences could be explained by CL-dependent deficiencies in SDH activity, which would also account for the observed differences in αKG levels (Fig. 5F).

ATP levels were reduced in Δ*crd1* mitochondria (Fig. 7A), consistent with previous reports that ATP synthesis is impeded in yeast Δ*crd1* mutants ^91^. The lowered starting concentration of ATP in Δ*crd1* mitochondria compared to both Δ*taz1* and WT mitochondria could be a consequence of ATP synthase hydrolyzing ATP to recoup the transmembrane potential in the face of a leaky membrane ^92^. The similar ATP levels for WT and Δ*taz1* in yeast were consistent with a recent report that found no difference in the rate of mitochondrial ATP production between healthy and BTHS patient iPSC-derived cardiomyocytes ^93^. Moreover, there is indirect evidence that defects in CL remodeling negatively influence both NAD^+ 94^ and NADH ^38^ levels, which appears to be consistent with our observations in Figure 7.

### BCAA, butanediol, and metabolic pathways unique to yeast mitochondria

Only ∼1% of the initial ^13^C pyruvate label was observed in the TCA cycle, with the majority distributed in BCAA pathways that use acetolactate or citramalate as entry points (Fig. 4). This is not unexpected given that the concentrations of intermediates in a self-contained cyclic pathway will remain low unless specific bottlenecks occur. In contrast, for cataplerotic pathways, intermediates that require cytosolic enzymes for further biosynthesis will accumulate over time in samples of isolated mitochondria. Variation in the kinetic profiles of metabolites such as citramalate (Fig. 5C), acetoin (Fig. 5D), acetolactate (Fig. 5I), αKI (Fig. 5J), and 2IPM (Fig. 5K) suggest there are differences in the yeast specific BCAA, citramalate, and butanediol pathways between the CL biosynthesis and remodeling mutant mitochondria and WT mitochondria. However, in contrast to the TCA cycle metabolites, none of the metabolites observed in the BCAA or butanediol pathway are reported to be synthesized by proteins associated with the IMM; therefore, any effect of IMM CL content on the function of these pathways could either be indirect and/or related to defective metabolite flux across the IMM.

### Mevalonate synthesis as a reporter for β-hydroxy β-methylglutaryl coenzyme A (HMG-CoA)

A particularly interesting profile is that for mevalonate (Fig. 5C), which showed elevated levels in Δ*taz1* compared to Δ*crd1* and WT. While mammals synthesize sterols in the endoplasmic reticulum and cytosol, some organisms like yeast ^81^ and trypanosomes ^95^ have mitochondrial HMG-CoA reductase enzymes that can make mevalonate in mitochondria. In yeast, mevalonate is synthesized from 3-hydroxy-3methylglutaryl-CoA (HMG-CoA) by HMG-CoA reductase ^81^.

A hallmark trait of BTHS patients is the presence of 3-methylglutaconic acid (3MGA) in their urine. A link between “primary” 3MGA-uria and enzymes involved in leucine degradation has been established ^96^. However, for “secondary” 3MGA-uria in diseases that include BTHS, there are no defects in leucine metabolism, and the origin of 3MGA is unknown ^23,96^. It has been proposed that 3MGA could be formed under conditions of decreased TCA flux (such as those observed for the CL biosynthesis and remodeling mutants here) when acetoacetyl-CoA synthase 2 converts acetoacetyl-CoA and acetyl-CoA to HMG-CoA, followed by hydrolysis of excess product to form CoA and 3MGA ^23,96^. It was further proposed that the alternative mitochondrial synthesis pathway could be confirmed by looking for 3MGA in isolated mitochondria ^96^. While we do not see 3MGA or HMG-CoA in our spectra, we do see mevalonate, which is the immediate product of HMG-CoA under the action of HMG-CoA reductase in yeast (Fig. 4).

The levels of mevalonate detected in mitochondria isolated from the Δ*taz1* yeast strain were more than three-fold larger than those from Δ*crd1* or WT (Fig. 5C). The Δ*taz1* mitochondria lack the phospholipid transacylase tafazzin, whose loss-of-function mutations cause BTHS and elevated 3MGA levels ^96^. The higher levels of mevalonate in the Δ*taz1* strain imply a higher level of the HMG-CoA precursor (Fig. 4). Moreover, it is difficult to rationalize higher levels of HMG-CoA with a purely vacuole origin of this metabolite in yeast, since tafazzin is located exclusively in the mitochondrion ^97^. The incorporation of the ^13^C-label into mevalonate, together with the higher levels of the metabolite in the Δ*taz1* strain, suggest that 3MGA and its precursor HMG-CoA can be synthesized *de novo* in mitochondria, in support of a previously proposed mechanism for 3MGA production in mitochondria ^96^.

### Yeast as a model system for studying CL biosynthesis and remodeling

Taken together, the results of the present study confirm the hypothesis that perturbations in the CL content of mitochondrial membranes strongly impacts metabolism. We observed differences between WT mitochondria and mitochondria from strains defective in CL biosynthesis and remodeling in the kinetic profiles of mevalonate, αKG, pyrimidine (NAD^+^/NADH) and adenine (ATP/AMP) nucleotides, and BCAA synthesis intermediates unique to yeast. Thus, the yeast system developed in this work could be further used to address additional basic biological questions in mitochondrial metabolism.

The use of yeast as a model system to investigate the molecular mechanisms of human disease is complicated by the large evolutionary distance between yeast and humans. This is highlighted by the fact that the bulk of the ^13^C_3_-pyruvate substrate in our study was metabolized by BCAA synthesis pathways that do not exist in humans. Therefore, these pathways unique to yeast confound direct comparisons in metabolism between the two organisms. Despite this drawback, the *in-organello* NMR metabolomics approach described herein should be translatable to mammalian tissue and cell culture models. As a result, we are currently applying it to more relevant models of human disease and seek to use it as a platform to screen for drugs that may remedy defects in mitochondrial pyruvate metabolism.

## EXPERIMENTAL PROCEDURES

### Chemicals and Reagents

Uniformly labeled ^13^C_3_-pyruvate and sodium-3-trimethylsilylpropionate (2,2,3,3-d_4_) (TSP) were from Cambridge Isotope Laboratories (Tewksbury, MA). All other chemicals and antibodies, including the mitochondrial pyruvate carrier inhibitor UK5099, were from Sigma-Aldrich.

### Preparation of Isolated Mitochondria

*Saccharomyces cerevisiae* (Brewer’s Yeast) mutant strains used for this work are isogenic to GA74-1A and were created as described previously ^35^. Yeast cells were grown at 30°C to mid-log phase (as determined by absorbance at 600 nm, A_600nm_ ∼1.2) in rich lactate medium (1% yeast extract, 2% tryptone, 0.05% glucose, 2% lactic acid, 3.4 mM CaCl_2_, 8.5 mM NaCl, 2.95 mM MgCl_2_, 7.35 mM KH_2_PO_4_, 18.7mM NH_4_Cl). Mitochondria were isolated by Dounce homogenization and differential centrifugation as described previously ^98^. Purified mitochondria were resuspended in SEH buffer (600 mM sorbitol, 1 mM Ethylenediaminetetraacetic acid (EDTA), 20 mM N-2-hydroxyethylpiperazine-N’-2-ethanesulfonic acid (HEPES), 0.5 mg/ml Bovine Serum Albumin (BSA), pH 7.5) containing protease inhibitors (1 mM phenylmethylsulfonyl fluoride (PMSF), 1 μg/ml aprotinin, 1 μg/ml leupeptin). Samples were divided into aliquots corresponding to 1 mg of mitochondrial protein, snap frozen in liquid nitrogen, and stored at -80°C until NMR measurements.

### NMR spectroscopy

Once-frozen mitochondria were thawed on ice for 30 minutes and sedimented for 5 minutes at 20,000xg and 4°C. The pelleted mitochondria were prepared similarly to a published protocol ^54^. They were first resuspended in 1 ml of PBS (137 mM NaCl, 2.7 mM KCl, 10 mM Na_2_HPO_4_, 1.8 mM KH_2_PO_4_, pH7.4) and re-spun at 20,000xg for 5 minutes, followed by resuspension in 200 μl of NMR analysis buffer (120 mM KCl, 5 mM KH_2_PO_4_, 1 mM EGTA, and 3 mM HEPES, pH 7.4). After incubating the isolated mitochondria on ice for 20 minutes ^36,54^, 400 µl of the respiratory substrate mix (3 mM ^13^C_3_-pyruvate, 0.25 mM malate, 0.25 mM NADH, 0.1 mM ADP, 0.1 mM ATP) were added to give a final NMR sample volume of 665 µl. D_2_O was added to a final concentration of 10% v/v for the NMR deuterium lock, and the NMR chemical shift reference and intensity standard TSP (trimethylsilylpropanoic acid) was included to a final concentration of 3 mM ^54^.

Kinetic profiles of metabolites were monitored with a total of 15 ^1^H-^13^C HSQC experiments, each collected in ∼5.5 minutes over a total experiment period of ∼90 minutes, with the addition of the ^13^C_3_-pyruvate substrate defined as the starting point of the reactions (t=0). For experiments looking at the inhibition of mitochondrial pyruvate transport, the UK5099 pyruvate carrier inhibitor was included at a final concentration of 3 mM. NMR experiments were performed on Varian INOVA 600 MHz and Bruker AVANCE 500 MHz spectrometers at a temperature of 30^°^C. For additional 1D NMR experiments monitoring downfield NMR signals,all experimental conditions were the same except^12^C_3_-pyruvate was used instead of ^13^C_3_-pyruvate.The 1D NMR experiments were obtained using the 1D-NOE pulse sequence *tnnoesy* on a Varian 600 MHz NMR instrument equipped with a cryogenic probe. Spectra were processed with iNMR (Mestrelab Research, Santiago de Compostela, Spain) and peak heights were measured with CcpNmr Analysis version 2.5.2 ^99^. Metabolite concentrations in units of fmol of metabolite per ng of mitochondrial protein were calculated relative to the peak height of 3 mM internal TSP standard.

### NMR Assignment of Metabolites

Assignments of metabolite NMR signals (summarized in Table S1) were based on matching ^1^H and ^13^C chemical shifts to known reference compounds in metabolomics databases including BMRB ^44^, PubChem ^42^, SpectraBase ^45^, SpinAssign ^41^, COLMAR ^47^, HMDB ^46^, and YMDB ^43^. In determining NMR assignments, we paid particular attention to whether the compound is found in yeast, and whether it is present in mitochondria (see references in Table 1). NMR assignments were further verified using ‘spiking’ experiments for the metabolites pyruvate, mevalonate, glutamate, glutamine, citrate, and succinate. The compounds acetolactate and acetoin that are unique to yeast ^76,78,100^ proved particularly difficult to assign since they are not well represented in metabolomics databases. Since these two metabolites gave the strongest signals in our HSQC experiments, they were present at a high enough concentration to observe in ^13^C HMBC experiments. Their assignments were thus enabled through long-range couplings observed in extracted metabolites from the mitochondria samples that made it possible to identify ^13^C ketone signals at 215 ppm (acetolactate) and 219 ppm (acetoin) (Fig. S3).

### Respirometry

Oxygen consumption rates of isolated mitochondria were measured using an O2k-FluoRespirometer (Oroboros Instruments, Innsbruck, Austria). After performing background calibration following the manufacturer’s instructions, 1 mg of mitochondrial protein (as determined by Bradford assay) was thawed on ice for 30 minutes, spun at 20,000xg for 5 minutes at 4°C, and resuspended to 4 mg/ml in Yeast Respiratory Media (225 mM sucrose, 75 mM mannitol, 10 mM Tris-HCl, 10 mM KH_2_PO_4_, 5 mM MgCl_2_, and 10 mM KCl) pre-warmed to 30°C. 60 µg of mitochondrial protein were added to each chamber, and once the O_2_ levels stabilized, pyruvate, malate, and NADH were added to final concentrations of 3 mM, 125 µM, and 125 µM, respectively to induce state 2 (basal) respiration. After reaching steady-state 2 respiration (∼11 minutes), ADP was added to a final concentration of 250 µM to reach steady-state 3 respiration. As the mitochondria consumed ADP to produce ATP, they transitioned to state 4 respiration; once reaching steady-state 4 respiration, mitochondrial oxygen consumption was terminated by adding antimycin A to a final concentration of 2.5 µM. Respiration rates were calculated by taking the slopes of the corresponding linear portions of O_2_ levels as a function of time. For pre-incubation measurements, mitochondrial respiratory activity was measured immediately after resuspension in Yeast Respiratory Media. For post-incubation measurements, resuspended mitochondria were incubated at 30°C for 65 minutes prior to assessment of oxygen consumption activity.

### Western Blot Analysis

Mitochondria were solubilized in SDS-PAGE sample buffer containing DTT, and mitochondrial proteins were separated using 12% gels. Proteins were transferred to PVDF membranes using an eBlot (GenScript Biotech, Piscataway, NJ) and the membranes were Western blotted for Tom40 and Tom70. Quantification of band intensity was performed using ImageJ ^101^.

### Statistics, rigor, and reproducibility

All reported means were obtained by averaging n ≥ 3 independent measurements across mitochondria obtained from three separate isolation procedures. Error bars represent means ± standard error of the mean (SEM) unless otherwise noted. Statistical tests were performed using one-way ANOVA and Tukey’s post-hoc analyses, where appropriate. Statistical significance was defined as p < 0.05.

## CONTRIBUTIONS

A.J.R. designed and performed experiments, analyzed the data, wrote the initial draft, and revised the manuscript. W.M. designed and performed the initial experiments, analyzed the data, wrote the initial draft, and revised the manuscript. S.M.C revised the manuscript and analyzed the data. N.N.A. advised A.J.R. and W.M., designed the experiments, revised the manuscript, analyzed the data, and provided funding. A.T.A. designed and performed experiments, advised A.J.R. and W.M., analyzed the data, wrote the initial draft, and revised the manuscript.

## ACKNOWLEDGEMENTS

This work was supported by the National Institutes of Health grant R01-AG065879.

## Notes

### Competing Interest Statement

The authors have declared no competing interest.

