## Supporting Materials for "Perturbations in mitochondrial metabolism associated with defective cardiolipin biosynthesis: An *in-organello* real-time NMR study"

**Table of contents:**

**Table S1 –** Metabolite YMDB accession codes, chemical shifts, and structures

**Supplementary Figure S1** – Uncropped Tom70 Western blots

**Supplementary Figure S2** – Mitochondrial respirometry

**Supplementary Figure S3** – NMR correlations used to verify metabolite assignments

**Table S1: Metabolite YMDB accession codes, chemical shifts, and structures**

| Metabolite | YMDB ID | Peak Number | $^{1}H$ Shift (ppm) | $^{13}C$ Shift (ppm) | C Atom | Structure |
|---|---|---|---|---|---|---|
| Pyruvate | 175 | 1 | 2.37 | 29.37 | 1 | 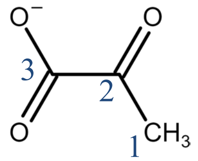 |
| Pyruvate Tautomer | | 1t | 1.49 | 28.08 | 1 | 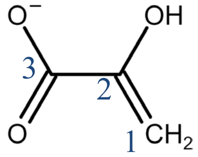 |
| Mevalonate | 707 | 2 | 1.34 | 28.05 | 6 | 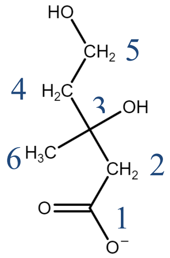 |
| Citramalate | 1584 | 3 | 1.23 | 27.23 | 5 | 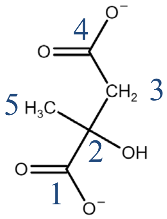 |
| 2-isopropylmalate | 1487 | 4<br>7<br>11 | 0.91<br>0.86<br>1.87 | 18.73<br>20.13<br>37.70 | 6<br>6’<br>5 | 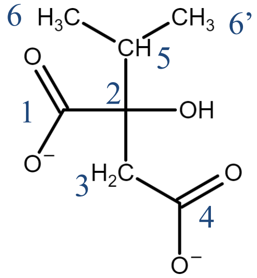 |
| Acetoin | 410 | 5 | 1.36 | 19.32 | 3 | 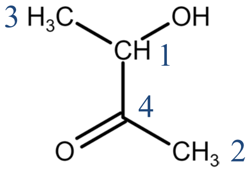 |

| | | | | | | |
|---|---|---|---|---|---|---|
| Valine | 152 | 6<br>8 | 0.95<br>1.00 | 19.19<br>19.52 | 4<br>4’ | 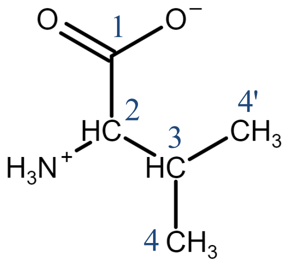 |
| Acetate | 59 | 9 | 1.93 | 26.13 | 2 | 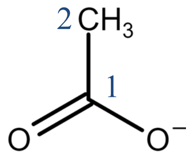 |
| Glutamate | 271 | 10 | 2.34 | 35.41 | 4 | 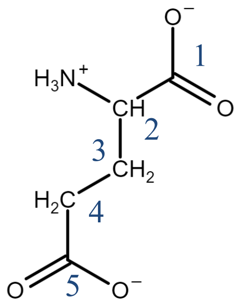 |
| Citrate | 86 | 12 | 2.68 | 47.89 | 1,3 | 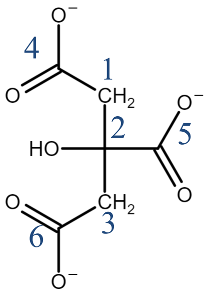 |
| α-ketoisovalerate | 365 | 13<br>15 | 2.94<br>1.13 | 39.85<br>19.23 | 1<br>2,2’ | 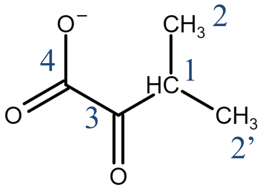 |

| | | | | | | |
|---|---|---|---|---|---|---|
| α-ketoglutarate | 2 | 14 | 2.44 | 33.51 | 4 | 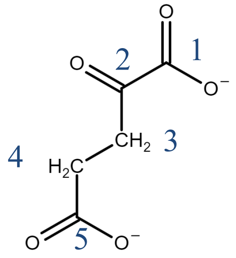 |
| Acetolactate | 394 | 16<br>17 | 1.47<br>2.26 | 24.20<br>27.27 | 3<br>1 | 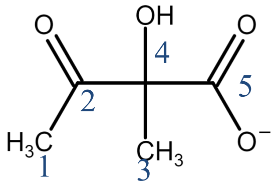 |
| Succinate | 338 | 18 | 2.41 | 37.36 | 2, 3 | 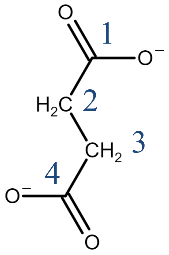 |

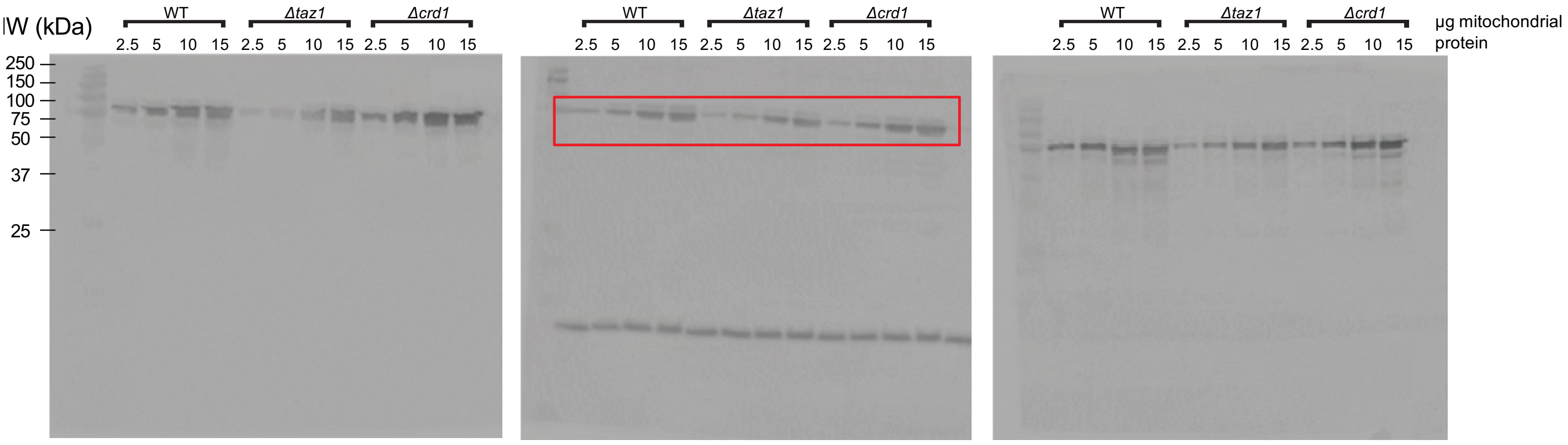


**Supplementary Figure S1: Uncropped Tom70 western blots**

Uncropped Western blots of Tom70, used for quantification in Figure 2B. Red box indicates cropped image used as representative blot in Figure 2A.

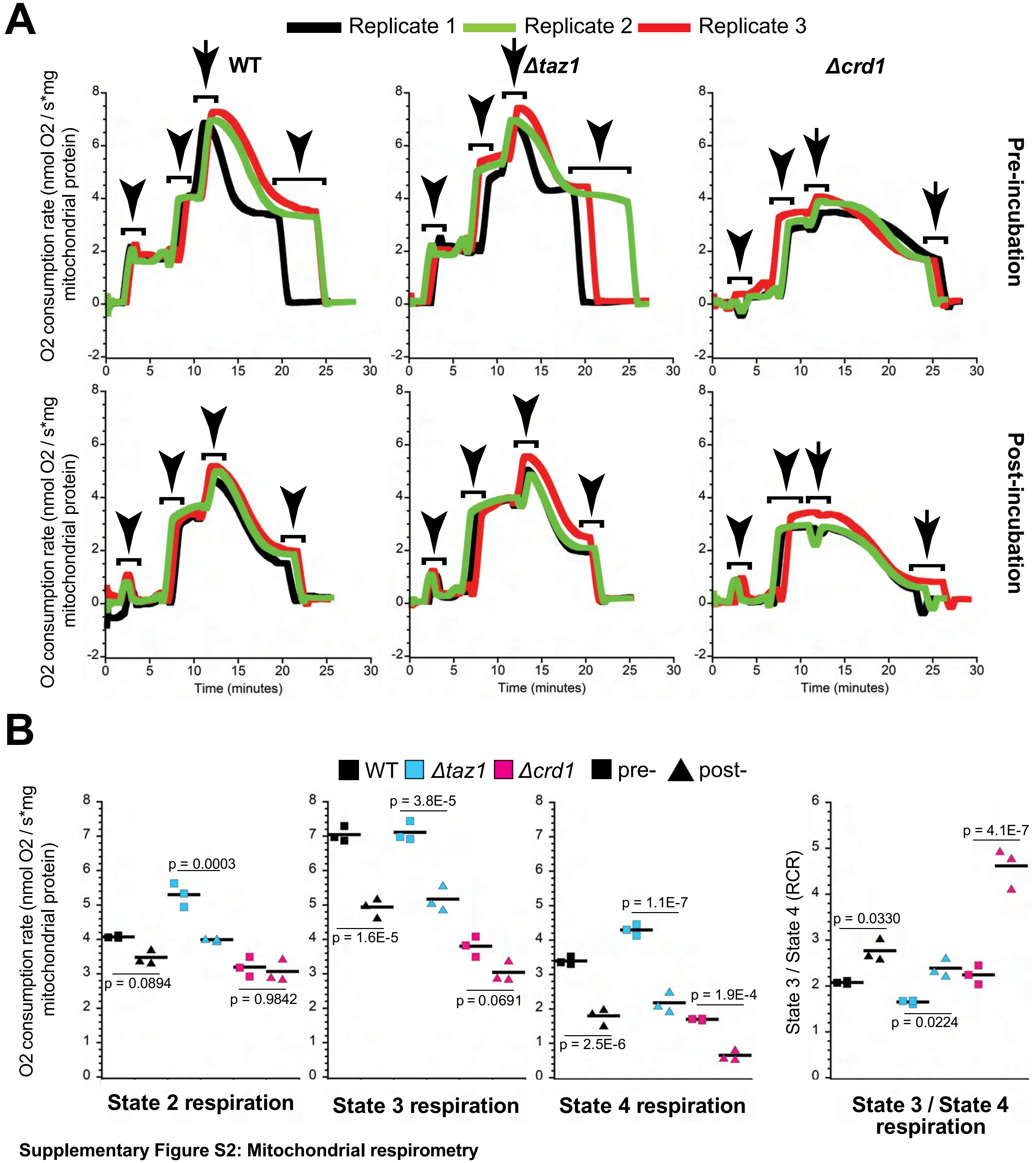


**Supplementary Figure S2: Mitochondrial respirometry**

*(A) Traces from 3 independent biological replicates pre- (top) and post- (bottom) 65 minutes of incubation at 30°C.* Arrows represent sequential additions of mitochondria (first arrow), respiratory substrates (malate, pyruvate, and NADH; second arrow), ADP (third arrow), and antimycin A (fourth arrow). *(B) Quantification of respiration rates pre- and post-65 minutes of incubation at 30°C.* State 2 respiration rates were obtained following addition of respiratory substrate (malate, pyruvate, and NADH), State 3 respiration rates were obtained following addition of ADP, State 4 respiration rates were obtained as respiration rates stabilized to a new steady-state after ADP phosphorylation, and respiration was stopped by the administration of antimycin A. p-values were determined by one-way ANOVA and Tukey's post-hoc test.

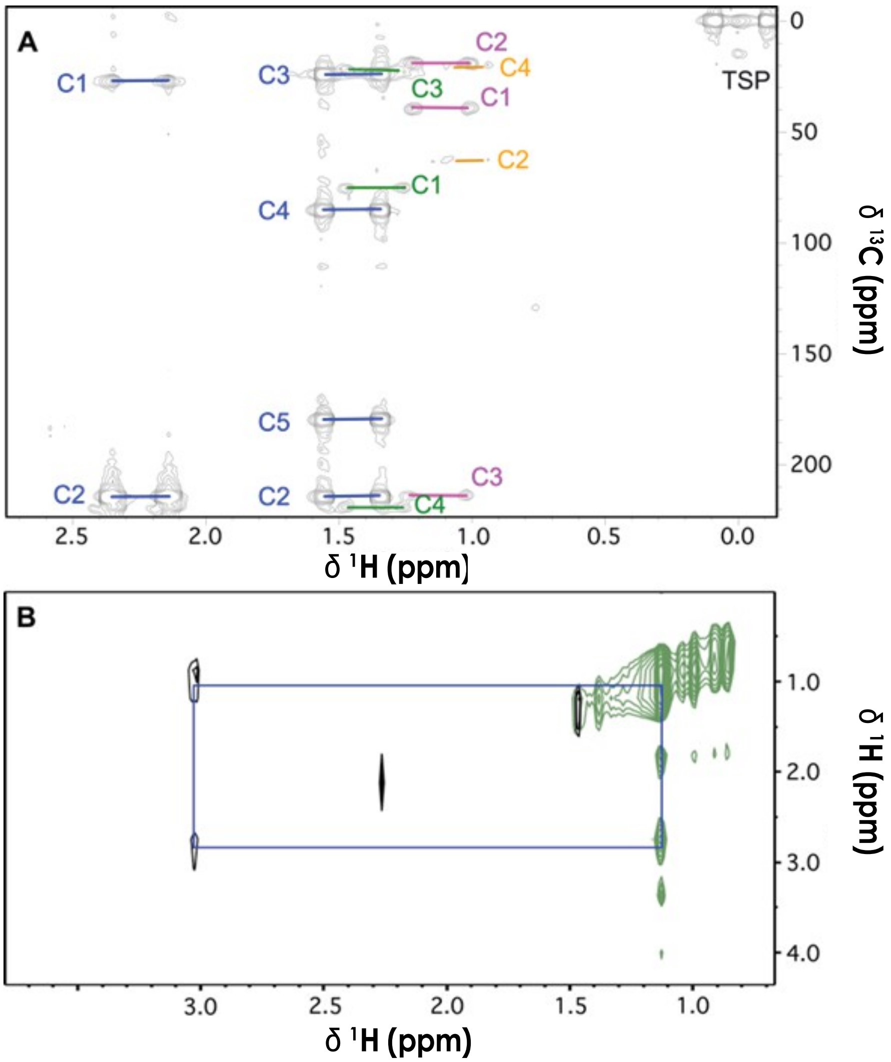


**Supplementary Figure S3: NMR correlations used to verify metabolite assignments**
*(A) HMBC spectrum showing long-range couplings that establish carbon frameworks for acetolactate (blue), acetoin (green), αKI (pink), and valine (orange).* Please refer to Table S1 for the carbon atom numbering. *(B) Superposition of $^{13}C$ planes from a 3D $^{13}C$ HSQC-TOCSY experiment: 19.1 ppm, green; 41 ppm, black.* The symmetric TOCSY cross peaks in both of the superposed planes connect the αKI C1 and C2 carbons (Table S1) and indicate both sites have $^{13}C$-label incorporated.
